# OmicsPharLeuDB: an integrative database for mining pharmacogenomic data in acute lymphoblastic leukemia

**DOI:** 10.1101/2023.09.14.557519

**Authors:** Yu Zhong Peng

## Abstract

Leukemia is one of the most common hematological malignancies in children. Recent cancer pharmacogenomic studies profiled large panels of cell lines against hundreds of approved drugs and experimental chemical compounds. However, until now, no comprehensive databases have been specifically designed for leukemia. Therefore, we have developed OmicsPharLeuDB (mugpeng.shinyapps.io/leu_web_english_peng_v2/), a database that includes the largest published studies investigating the viability response of human leukemia cell lines to chemical compound treatment, and the association between drug sensitivity and multi-omics, and also provides a user-friendly interactive website allowing the user to assess drug sensitivity data from a different platform and find the potential relationship between drugs and omics.

## Introduction

The emergence of high-throughput technologies has led to the generation of a vast amount of genomic data across various disease domains. In oncology, the genomic and pharmacological profiling of cancer cell line models has significantly improved our understanding of the relationship between the molecular features of cancers and treatment outcomes. Multiple studies have produced pharmacogenomic profiles and multi-omics data for cancer cell lines, such as the Genomics of Drug Sensitivity in Cancer (GDSC) and the Cancer Therapeutic Response Portal (CTRP). Several databases have been developed to integrate these cancer cell line pharmacogenomic datasets, including PharmacoDB [1], iGMDR [2], and DrugComb [3]. However, none of these databases are specifically designed for leukemia and provide limited insights into the association between multi-omics results and drug sensitivity. Therefore, we have designed and developed a user-friendly, comprehensive database known as multi-omics, pharmaceutics related leukemias database “OmicsPharLeuDB” (mugpeng.shinyapps.io/leu_web_english_peng_v2/), which integrates genomic and pharmacological profiles data collected from seven different databases, involving seven lineage subtypes, and containing a total of seven omics data. It also has several functionalities which allows user to find drugs with the highest drug sensitivity and explore the correlation between an omic like gene expression profiles and drug sensitivity.

## Methodology and implementation

### Data collection

We collected data from several published cancer or leukemia studies: the Cancer Therapeutic Response Portal (CTRP1,2) [4,5], Genomics of Drug Sensitivity in Cancer (GDSC), Genentech Cell Screening Initiative (gCSI) [6], BeatAML [7], the work of Tavor et al. [8], and Functional Omics Resource of ALL(FORALL) [9]. These datasets included drug sensitivity data and high through-out omics data of human cancer cell lines. We subseted the leukemia data if the dataset compromising other cancer types. As a result, we got 528 cells and 123 drugs from beatAML, 9 cells and 342 drugs from CTRP1, 71 cells and 482 drugs from CTRP2, 43 cells and 529 drugs from FORALL, 82 cells and 305 drugs from GDSC1, 70 cells and 153 drugs from GDSC2, and 50 cells and 47 drugs from Tavor’s work respectively (Fig 1A), and seven leukemia sub-lineage types including Acute Lymphoblastic Leukemia(ALL), Acute Biphenotypic Leukemia(ABL), Acute Myelogenous Leukemia(AML), Chronic Lymphoblastic Leukemia(CLL), Chronic Myeloid Leukaemia(CML), Hairy Cell Leukemia(HCL) and Acute Promyelocytic Leukemia(APL). The omics data we collected including mRNA expression, protein data, copy number variant(CNV), gene fusion, methylation, mutation(gene mutation, and a specific amino acid change). The details of collected datasets including the numbers of cell lines and drugs and omics information were listed in Table 1.

**Figure 1.**
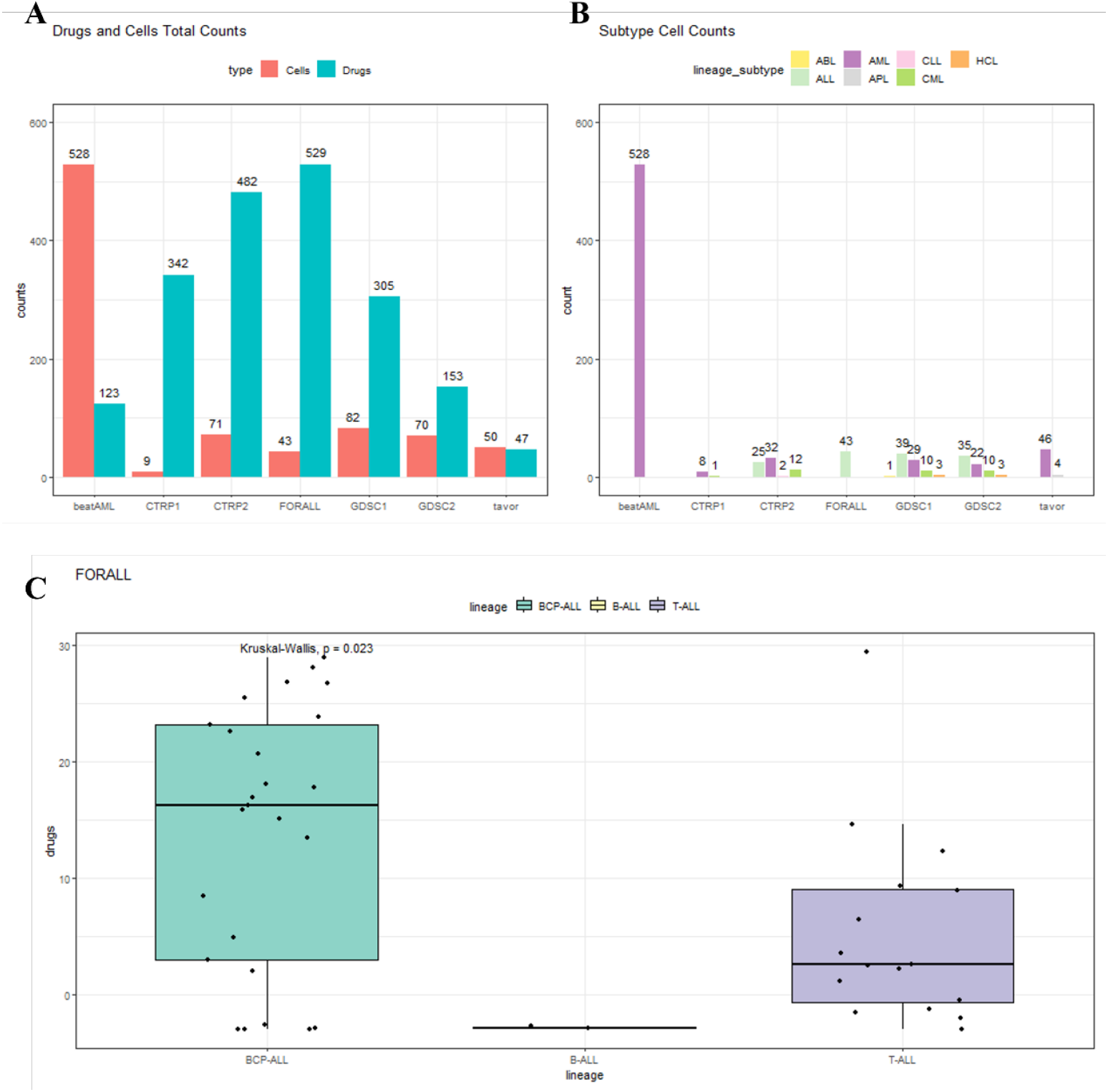
Contents of ‘OmicsPharLeuDB’. Total number of (A) drugs, cells and (B) subtypes in seven collected projects. (C) Drugs’ sensitivity in different sub-lineage.

**Figure 2.**
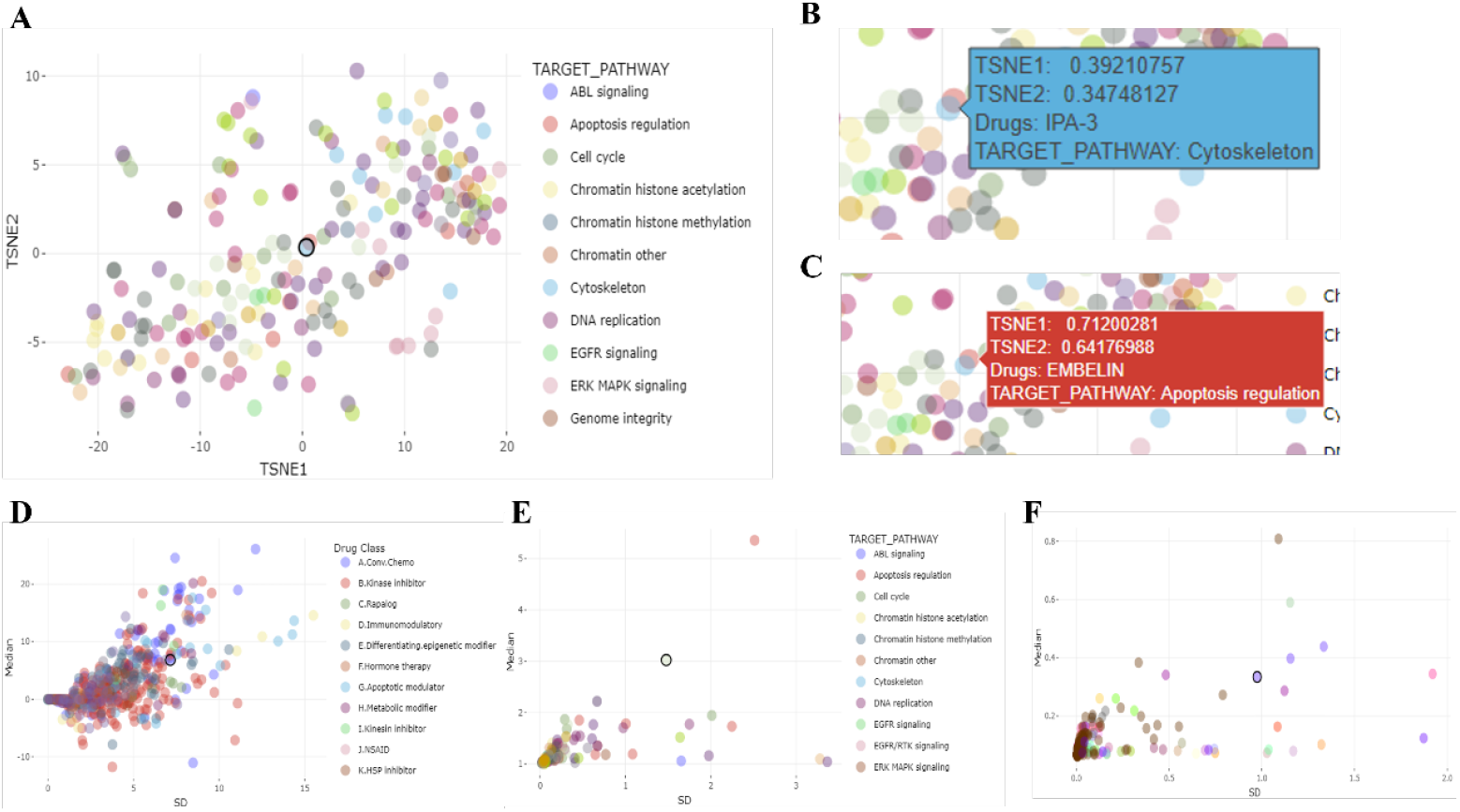
Scatter plot of profile of drug sensitivity, drugs are highlighted and colored according to their drug class. (A) TSNE of the distribution of drugs as a two-dimensional reduction in GDSC1. Drug sensitivity metrics were used to generate the UMAP. (B,C) UMAP coordinates of IPA-3 and EMBELIN. The median versus variance for each drug across the cell lines in FORALL (D), CTRP1(E), GDSC2(F).

**Table 1.**
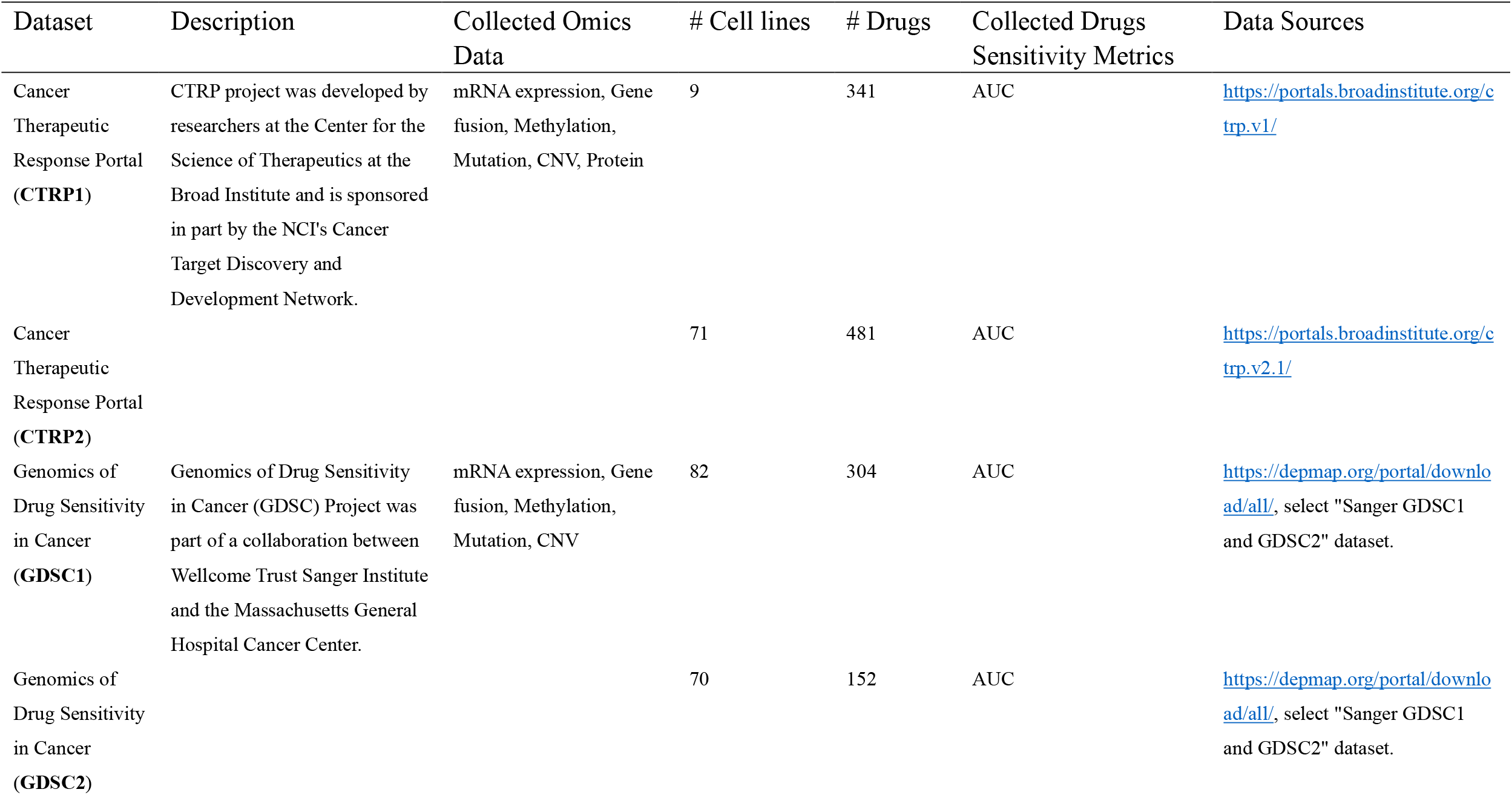
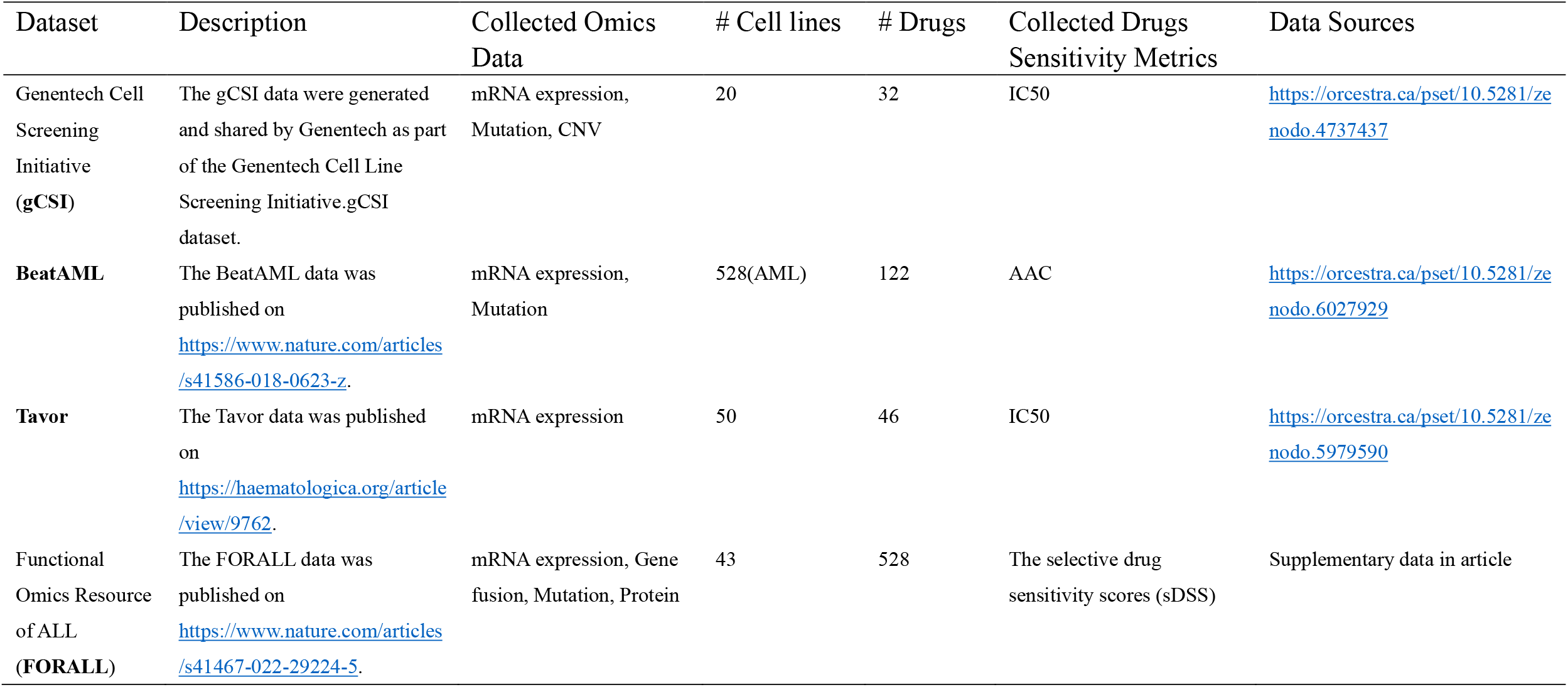
Details of datasets in OmicsPharLeuDB.

| Dataset | Description | Collected Omics Data | # Cell lines | # Drugs | Collected Drugs Sensitivity Metrics | Data Sources |
| --- | --- | --- | --- | --- | --- | --- |
| Cancer Therapeutic Response Portal (CTRP1) | CTRP project was developed by researchers at the Center for the Science of Therapeutics at the Broad Institute and is sponsored in part by the NCI's Cancer Target Discovery and Development Network. | mRNA expression, Gene fusion, Methylation, Mutation, CNV, Protein | 9 | 341 | AUC | https://portals.broadinstitute.org/ctrp.v1/ |
| Cancer Therapeutic Response Portal (CTRP2) |  |  | 71 | 481 | AUC | https://portals.broadinstitute.org/ctrp.v2.1/ |
| Genomics of Drug Sensitivity in Cancer (GDSC1) | Genomics of Drug Sensitivity in Cancer (GDSC) Project was part of a collaboration between Wellcome Trust Sanger Institute and the Massachusetts General Hospital Cancer Center. | mRNA expression, Gene fusion, Methylation, Mutation, CNV | 82 | 304 | AUC | https://depmap.org/portal/download/all/, select "Sanger GDSC1 and GDSC2" dataset. |
| Genomics of Drug Sensitivity in Cancer (GDSC2) |  |  | 70 | 152 | AUC | https://depmap.org/portal/download/all/, select "Sanger GDSC1 and GDSC2" dataset. |
| Genentech Cell Screening Initiative (gCSI) | The gCSI data were generated and shared by Genentech as part of the Genentech Cell Line Screening Initiative.gCSI dataset. | mRNA expression, Mutation, CNV | 20 | 32 | IC50 | https://orcestra.ca/pset/10.5281/zenodo.4737437 |
| BeatAML | The BeatAML data was published on https://www.nature.com/articles/s41586-018-0623-z. | mRNA expression, Mutation | 528(AML) | 122 | AAC | https://orcestra.ca/pset/10.5281/zenodo.6027929 |
| Tavor | The Tavor data was published on https://haematologica.org/article/view/9762. | mRNA expression | 50 | 46 | IC50 | https://orcestra.ca/pset/10.5281/zenodo.5979590 |
| Functional Omics Resource of ALL (FORALL) | The FORALL data was published on https://www.nature.com/articles/s41467-022-29224-5. | mRNA expression, Gene fusion, Mutation, Protein | 43 | 528 | The selective drug sensitivity scores (sDSS) | Supplementary data in article |

The drug sensitivity metrics including: AUC(area under the dose–response curve) (CTRP1,2 and GDSC1,2), IC50(relative half-maximal inhibitory concentration) (gCSI and Tavor), AAC(area above the dose–response curve) (BeatAML), and sDSS(selective drug sensitivity scores) (FORALL). Lower values of AUC and IC50 indicated increased sensitivity to drug treatment, while the higher area above the dose-response curve(AAC) and the selective drug sensitivity scores (sDSS) meant a higher treatment sensitivity.

### Data preprocess

As mentioned before, the database collected different metrics involving AUC, IC50, AAC and sDSS. In order to give the metrics a consistent direction and to represent stronger drug sensitivity with higher values, AUC and IC50 were convert into its reciprocal.

For the purpose of plotting T-SNE, the drugs with more than 20% missing records(NA) were discarded in each datasets. Then R package impute was used to embed a nearest neighbor averaging function to impute the remained missing data.

All preprocessed procedures were finished using R.

### Publication-ready plotting module

Plots are generated using R and basing on R packages, including ggplot2 [10], ggpubr [11], plotly [12], patchwork [13], paletteer [14], DT [15].

### Web server implementation

OmicsPharLeuDB web server was implemented on shinyapp(www.shinyapps.io) with Shiny R package [16]. Database is accessible from multiple platforms through Microsoft Edge, Chrome, Safari and Firefox.

## Utility and discussion

The database had three main parts: 1) Drugs sensitivity display; 2) Drugs-omics pairs analysis; 3) Features database significant analysis.

### Drugs sensitivity data display

This function consisted of two parts.

The first one was the comparison of a certain drug sensitivity between different cell subtypes. The user chose a certain drug, and it would return all datasets including this drug visualized as a boxplot with the significant test to check if there was an association between leukemia subtypes and a certain drug sensitivity metric. For example, the Prednisolone showed a significant drug sensitivity in BCP-ALL comparing with other cell lines from FORALL project (Fig1C).

The second part was the profile of drug sensitivity. The T-SNE dimensionality reduction plots were generated for comparing each drug for inspecting 1) if two drugs with similar drug targets but showing different drug sensitivity or 2) if two drugs with different drug targets but having close drug sensitivity. Besides, the median versus variance scatter would tell if a drug had a wide range sensitivity in different cell lines and its sensitivity rank in the database. For example, IPA-3 targeted cytoskeleton pathway showed a similar drug sensitivity like apoptosis regulation drug EMBELIN across cell lines in GDSC1 (Fig3A-C). Additionally, SD-MEDIAN plot indicated that VINCRISTINE was an effective drug both in CTRP1, GDSC2 and FORALL (Fig 3D-F), and provenly, it was a FDA-approved clinical drug for acute leukemia.

**Figure 3.**
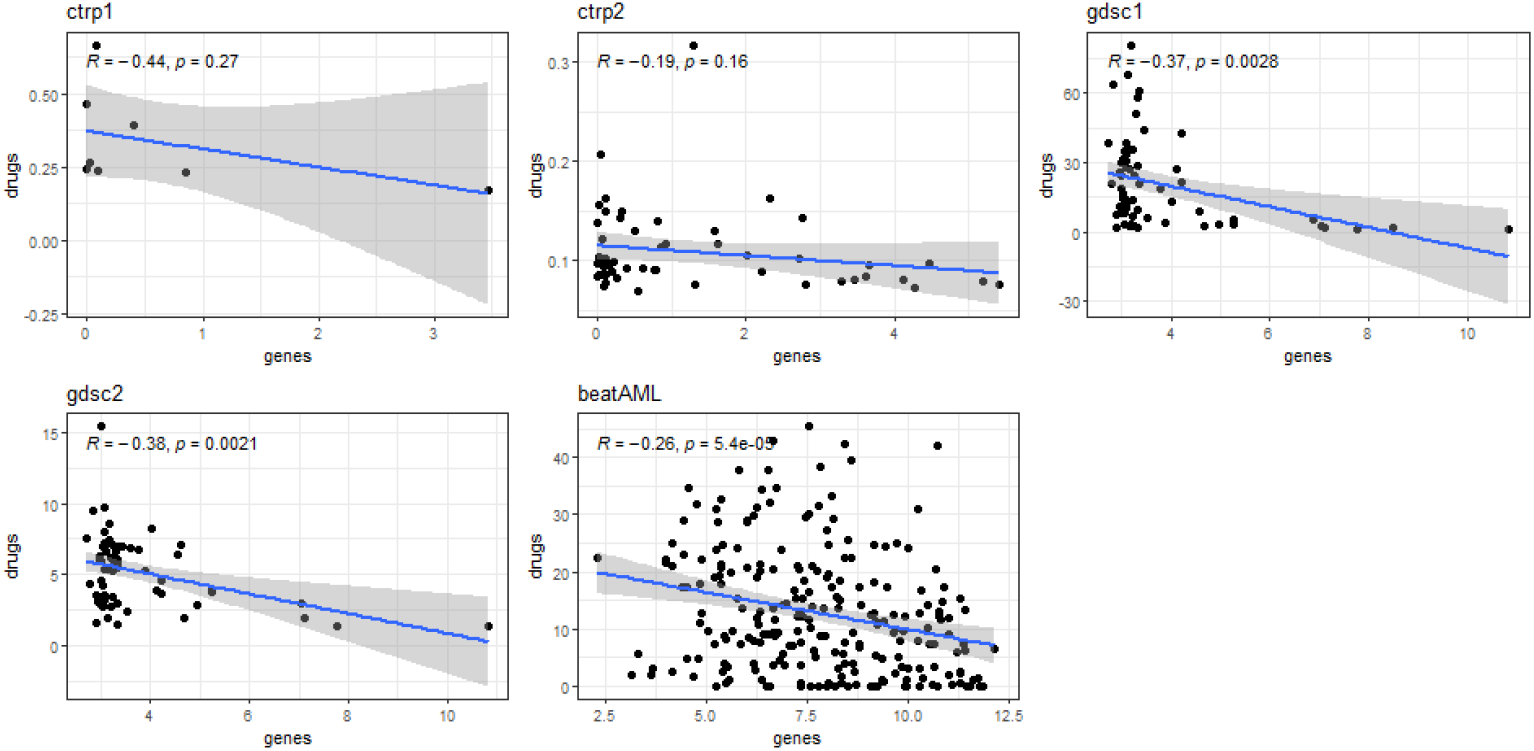
The correlation between the drug sensitivity summary metrics for the YM-155 and mRNA expression level of the ATP Binding Cassette Subfamily B Member 1, ABCB1, for the different ALL cell lines in five projects.

### Drugs-omics pairs analysis

This feature allowed user to explore the association between a selected drug resistance event and a certain omic. For continuous omics data like mRNA, methylation, copy number variant, protein, spearman correlation was calculated. While for discrete omics data such as mutation genes, mutation gene points or gene fusions, wilcoxon test was chosen for testing Signification. For example, the gene expression of ABCB1 was found to be negatively correlated with drug YM-155 in five datasets and three of them with P < 0.05 (Fig 3). Besides, the TP53 mutated group was identified to help drug resistance of SNX-2112 (Fig 4).

**Figure 4.**
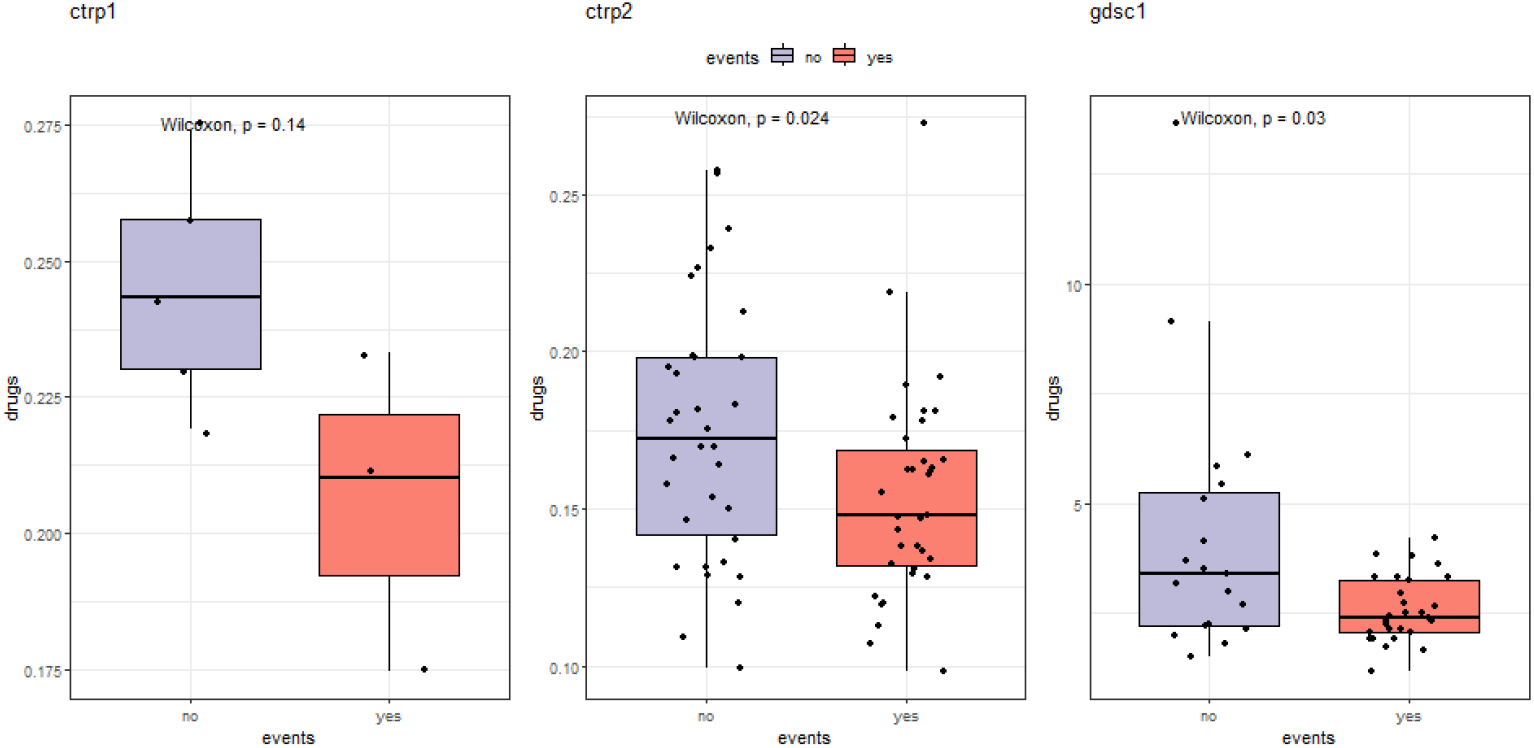
The drug sensitivity summary metrics of SNX-2112 in the TP-53 mutated cell lines compared to the wild type cell lines in different projects. The two-sided wilcoxon-test P value is indicated as (P)..

### Features database significant analysis

This analysis module helped people to conduct a significant test between a targeted feature(a drug or a omic) and all the features in a particular feature dataset grouped by their collected databases in a large scale. For example, the analysis returned the significant test results between mRNA expression of ABCB1 and all drug sensitivity metrics in six datasets, and showed that compound 1B was the highest drugs positively correlated with ABCB1 in ctrp2 dataset (Fig 5). The volcano plot included two axis, the y-axis is -log10 P value and the x-axis is the R/Diffs/Odds which compromising three conditions: 1) R(0-1) for continuous feature and continuous dataset, like drug A with all features in CNV; 2) Odds(>1 meant the selected feature A has more prop of feature B events from a certain database) for discrete feature and discrete database, like mutation events of TP53 and all collected gene fusion. And The features with q-value <= 0.1 are highlighted in blue. This step helped users to find all potential features with a certain interested omic or drug. Moreover, users was able to download the R list object used for visualization and did the subsequent analysis like filtering features, or collecting significantly correlated genes for enrichment analysis.

**Figure 5.**
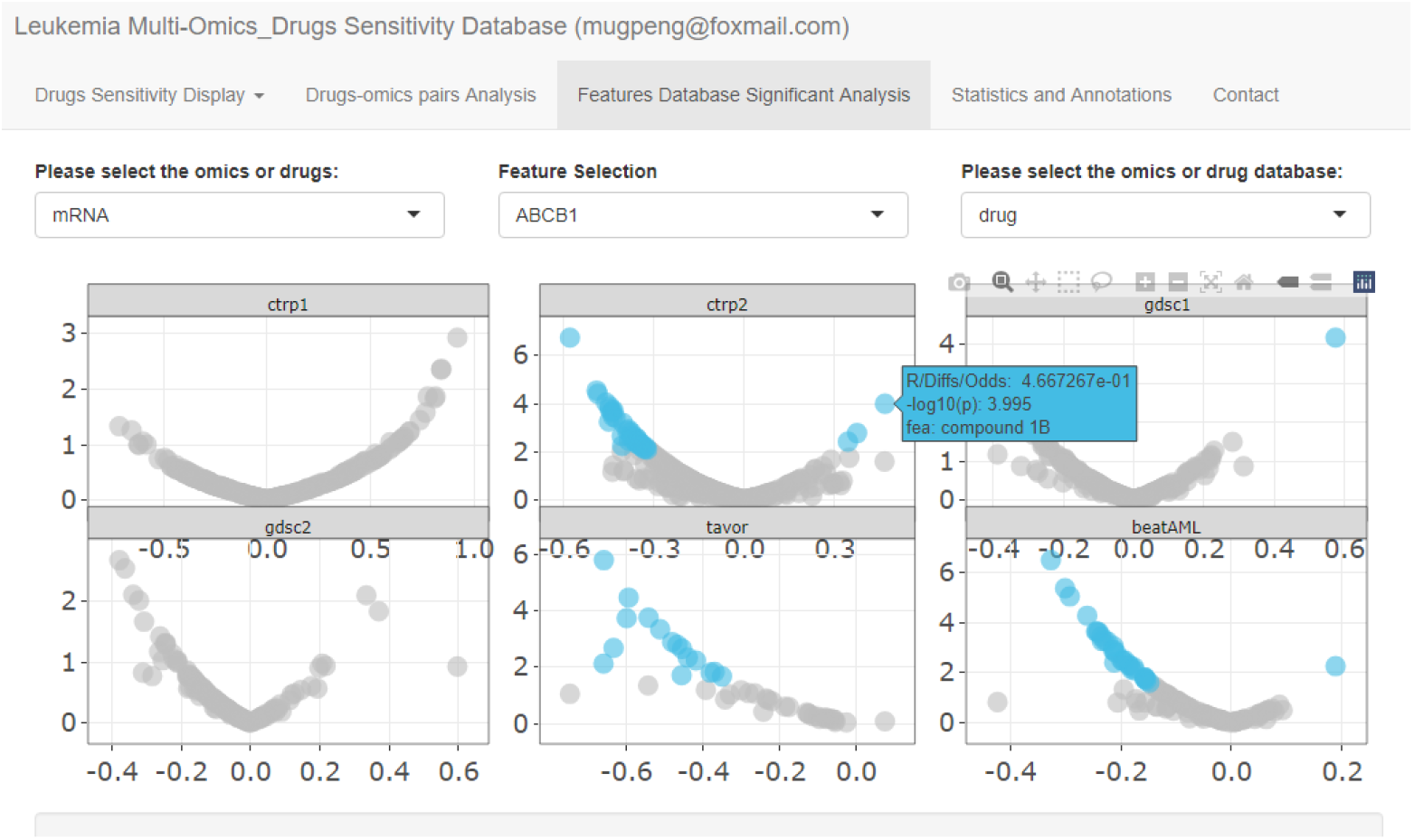
The interactive scatter plot to depict the correlation between selected feature and all data in an another feature database(mRNA, mutation, drug sensitivity, etc.).

## Conclusion

OmicsPharLeuDB was the first database providing a comprehensive resource to search and explored the largest pharmacogenomic studies specifically for ALL. By combining rigorous curation of identifiers across the published pharmacogenomic datasets with comprehensive search and visualizations of the pharmacological data, OmicsPharLeuDB allowed researchers to quickly access the data available to answer their biological questions of interest. It also provided an interface to do some analysis including drugs sensitivity display, drugs-omics pairs analysis and features database significant analysis.

## Availability of data and materials

The web server is available at https://mugpeng.shinyapps.io/leu_web_english_peng_v2. The source code for the web server and the standalone R package is available at github: https://github.com/mugpeng/OmicsPharLeuDB.

